# Rare Rewards Amplify Dopamine Learning Responses

**DOI:** 10.1101/851709

**Authors:** Kathryn M. Rothenhoefer, Tao Hong, Aydin Alikaya, William R. Stauffer

## Abstract

Dopamine neurons drive learning by coding reward prediction errors (RPEs), which are formalized as subtractions of predicted values from reward values. Subtractions accommodate point estimate predictions of value, such as the average value. However, point estimate predictions fail to capture many features of choice and learning behaviors. For instance, reaction times and learning rates consistently reflect higher moments of probability distributions. Here, we demonstrate that dopamine RPE responses code probability distributions. We presented monkeys with rewards that were drawn from the tails of normal and uniform reward size distributions to generate rare and common RPEs, respectively. Behavioral choices and pupil diameter measurements indicated that monkeys learned faster and registered greater arousal from rare RPEs, compared to common RPEs of identical magnitudes. Dopamine neuron recordings indicated that rare rewards amplified RPE responses. These results demonstrate that dopamine responses reflect probability distributions and suggest a neural mechanism for the amplified learning and enhanced arousal associated with rare events.

## Main Text

Making accurate predictions is evolutionarily adaptive. Accurate predictions enable individuals to be in the right place at the right time, choose the best options, and efficiently scale the vigor of responses. Dopamine neurons are crucial for building accurate reward predictions. Phasic dopamine responses code for reward prediction errors: the differences between the values of received and predicted rewards (*1-8*). These signals cause predictions to be modified through associative and extinction learning (*9, 10*). Likewise, phasic dopamine neuron stimulation during reward delivery increase both the dopamine responses to reward predicting cues and the choices for those same cues (*11*). Although it is well understood how predicted reward values affect dopamine responses, these predictions are simply point estimates – normally the average value – of probability distributions. It is unknown how the form of probability distributions affects dopamine learning signals.

Probability distributions affect behavioral measures of learning and decision making. Learning the expected value takes longer when rewards are sampled from broader distributions, compared to when they are drawn from narrower distributions (*12*). Likewise, even with well learned values, decision makers take a longer time to choose between options when the value difference between rewards is smaller, compared to when the difference is larger (*13, 14*). These behavioral measures of learning and decision making demonstrate that the probability distributions over reward values, and not simply the point estimates of values, influence learning and decision making. Prior electrophysiological recordings have demonstrated that dopamine signals adapt to the range between two outcomes (*15*), but it is not known whether these same dopamine signals reflect probability distributions over reward values.

Here, we asked whether the shapes of probability distributions were reflected in dopamine reward prediction error responses. We created two discrete reward size distributions that reflected, roughly, the shapes of normal and uniform distributions. We trained monkeys to predict rewards drawn from these distributions. Crucially, the normal distribution resulted in rare prediction errors following rewards drawn from the tails of that distribution, whereas the same rewards, with identical prediction errors, were drawn with greater frequency from the uniform distribution. We found that monkeys learned to choose the better option within fewer trials when rewards were drawn from normal distributions, compared to when rewards were drawn from uniform distributions. Moreover, we found that pupil diameter was correlated with rare prediction errors, but uncorrelated with common prediction errors of the same magnitude, suggesting greater vigilance to rare outcomes. Using single neuron recording, we show that dopamine responses reflect the shape of predicted reward distribution. Specifically, rare prediction errors evoked significantly larger responses than common prediction errors with identical magnitudes. Together, these results demonstrate a complementary but distinct mechanism from TD-like reward prediction error responses for learning based on probability distributions.

## Results

### Reward size distributions

We used non-informative images (fractal pictures) to predict rewards drawn from differently shaped distributions. Distribution shapes were defined according to relative reward frequency. One fractal image predicted an equal probability of receiving a small, medium, or large volume of juice reward. We define this as the uniform reward size distribution (Fig. 1A left). A second fractal image predicted that the small and the large reward would be given 2 out of 15 times, and the medium sized reward would be given the remaining 11 out of 15 times (normal reward size distribution, Fig. 1A right). Importantly, both reward size distributions were symmetrical and were comprised of the same three reward magnitudes. Therefore, the uniform and normal distributions had identical Pascalian expected values. However, rewards drawn from the tails of the normal distribution were rare, compared to the frequency of identical rewards drawn from the tails of the uniform distribution. Anticipatory licking reflected the expected value of both the distributions, as well as the expected value of safe cues (Figure 1B, *p* = 0.019, Linear Regression). Thus, the animals learned that the cues predicted rewards.

**Figure 1.**
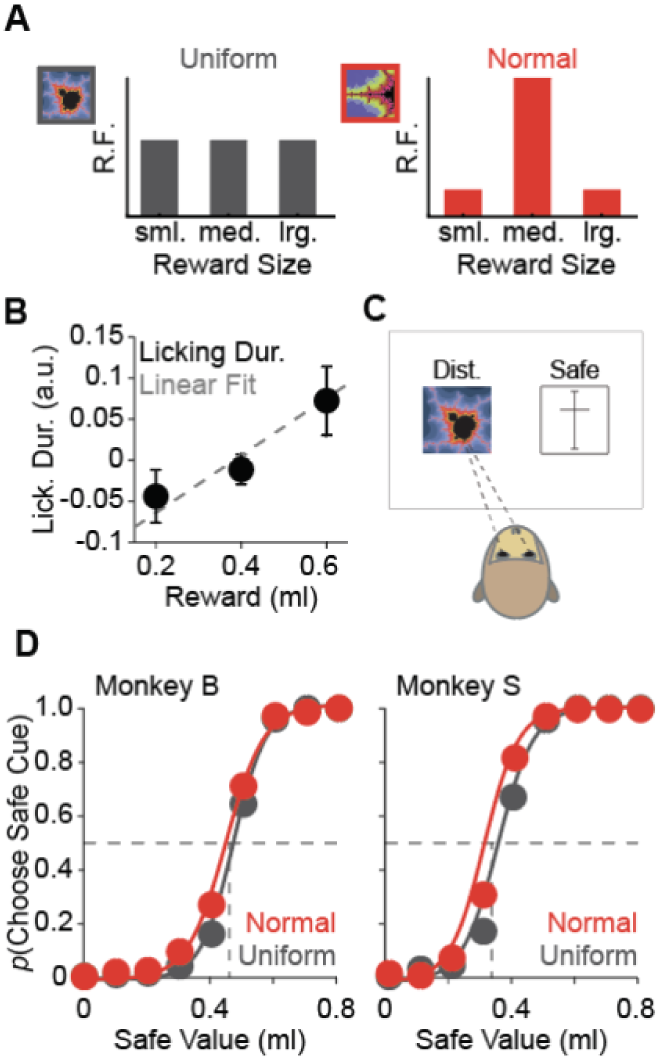
Reward size distributions. **A.** Fractal images predicted that one of three juice reward sizes would be drawn from a uniform (left) or normal (right) reward size distribution. The uniform distribution predicted that each size would be drawn with equal relative frequency (R.F.) (1/3 for small, medium, and large sized rewards). The normal distribution predicted that the small and large reward volumes would be drawn 2/15 times, and the medium reward volume would be drawn the remaining 11/15 times. **B.** Anticipatory licking indicated that the monkeys learned the predicted reward values. Black dots indicate the normalized licking duration data for predicted reward volume, and the grey dotted line indicates the linear fit to the data. Error bars indicate SEM across session. **C.** The choice task used to measure subjective value. Animals made saccade-directed choices between a distribution predicting cue and a safe alternative option. The safe alternative option was a value bar with a minimum and maximum of 0 and 0.8 ml at the bottom and top, respectively. The intersection between the horizontal bar and the scale indicated the volume of juice that would be received if monkeys selected the safe cue. **D.** Probability of choosing the safe cue as a function of the value of the safe option, when the distribution predicting cue had an expected value (EV) of 0.4ml. Dots show average choice probability for 9 safe value options for monkey B (left), and Monkey S (right). Solid lines are a logistic fit to the data. Red indicates data from normal distribution blocks, grey indicates data from uniform distribution blocks. The dashed horizontal lines indicate subjective equivalence, and the CE for each distribution type is indicated with the dashed vertical lines.

Dopamine neurons code prediction errors in the subjective values of rewards (*16*), according to the following equation:

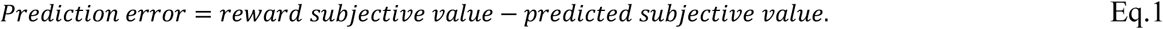

To investigate whether dopamine responses were sensitive to probability distributions, we sought to compare dopamine responses when the conventional prediction errors, defined according to Eq. 1, were identical. Therefore, we created reward distributions with the same expected values and then measured the distribution of the subjective values. We measured the subjective values of six reward distributions – three normal reward distributions with center locations of 0.3, 0.4, and 0.5 ml and three uniform distributions with center locations of 0.3, 0.4, and 0.5 ml. Monkeys made choices between cues that predicted a distribution and safe values (Fig. 1C, Methods). We plotted the probability of choosing the safe option as a function of the safe option volume and generated psychometric functions (Figure 1D). We used the certainty equivalents (CEs) – defined as the safe volume at the point of subjective equivalence between the two options (Fig. 1D, vertical dashed. Line) – as a measure of the distribution subjective value. Analysis of variance (ANOVA) on the CEs for the normal and uniform distribution cues indicated a significant effect of EV on CE (*F* = 71.81, *p* < 0.001, ANOVA). Crucially, we found no significant effect of the distribution type on the CEs (*F* = 3.57, *p* = 0.07, ANOVA). To increase our power to detect small differences in subjective value, we combined the CE data from all three EVs, yet still found no significant difference between the subjective values of the normal and uniform distributions (*p* = 0.20, two-sample *t*-test). Therefore, within the limited range of reward sizes we used, the data indicate that the normal and uniform reward size distributions had similar subjective values. These results indicated that the prediction errors generated from the distributions could be readily compared and ensured that disparities between prediction error responses were not driven by differences in the predicted subjective values.

### Distribution shape affects learning

We next investigated how probability distributions affected learning (Fig. 2A). Each learning block consisted of 15-25 trials and used two, never-before-seen cues (Supplementary Methods). Both cues predicted either normal or uniform distributions and had two different expected values (EVs). The animals had to learn the relative EVs to maximize reward. We analyzed behavioral choice data from two animals across 16 sessions, each of which included a mixture of normal and uniform blocks (Supplementary Methods). As expected, the probability of choosing the higher value option on the first trial of each block was not significantly different from chance (Fig. 2B, left). Comparison between the first and fifteenth trial of each block revealed that the animals learned to choose the higher value option on 87% and 85% of trials, in the normal and uniform blocks, respectively (Fig. 2B, *p* < 0.001, paired *t*-test for both distributions). There was no significant difference in the average performance between the fifteenth trials of normal and uniform blocks (Fig. 2B, right, *p* = 0.7, *t*-test). Together, these results demonstrate that the animals learned which option had a higher EV.

**Figure 2.**
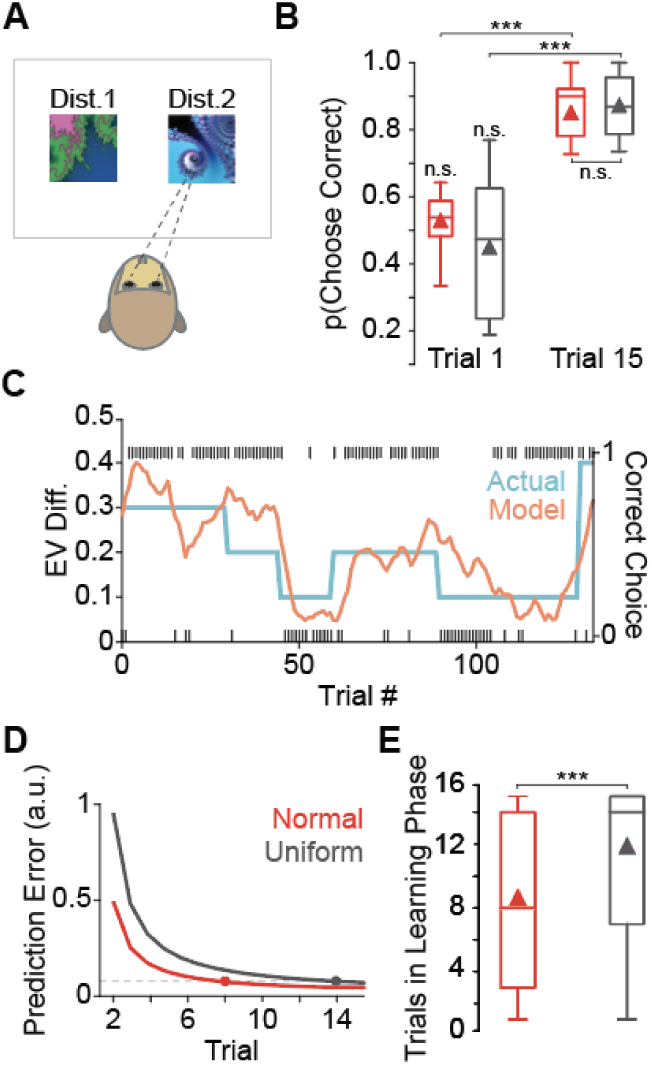
Faster learning from normal reward distributions. **A.** Animals made choices between two never-before-seen fractal images with different expected values (EVs), but the same distribution type (normal or uniform). **B.** Box and whisker plots show the probability of choosing the higher-valued option on trials 1 and 15, separated by normal (red) and uniform (grey) blocks. Triangles represent the averages. n.s. = not significantly different. **C.** Actual (blue) and estimated (orange) value differences for two choice options. The primary y-axis shows the EV differences between the two choice options, and the x-axis shows trial number. The block-wise changes in the true value difference are reflected by the model. The black tick marks correspond to correct and incorrect choices, defined by the relative expected values, and indicated by the secondary y-axis. **D.** Fitted prediction errors as a function of trial within normal (red) and uniform (grey) blocks. Dashed line indicated the threshold for designating that block values have been learned. Dots indicate the transition from the learning to the stable phase. **E.** Box and whisker plot showing the number of trials in the learning phase for normal and uniform distributions. The plot follows the same format as panel B.

We hypothesized that the animals would learn faster from reward sizes sampled from the normal distribution compared to the uniform distribution. We used a reinforcement learning (RL) model to quantify the learned values (Supplementary Methods). Our model, fit to the behavioral choices, performed well at predicting the true values of the two choice options (Fig 2C). To differentiate active learning from stable, asymptotic-like behavior, we fit a logarithmic function to the estimated prediction errors (Fig. 2D). When the change in the log-fitted values went below a robust threshold, we considered the values to be learned (Supplementary Methods). Using this approach, we determined the block-wise number of trials needed to learn. We found that, on average, animals needed 4 additional trials to learn in the uniform blocks, compared to the normal blocks (Fig. 2E, *p* < 0.001, Mann-Whitney U test). These data demonstrate that the monkeys learned faster from normal reward distributions, compared to uniform reward distribution. Thus, rare prediction errors have greater behavioral relevance than common prediction errors of the same magnitude.

To investigate autonomic responses to prediction errors, we analyzed the pupil responses during the choice task (Fig 3A). We used deconvolution to separate the effects of distinct trial events on pupil responses (Supplementary Methods). The deconvolution analysis indicated that the correlation between pupil responses and RPE was higher during the learning phase of normal blocks, compared to the learning phase of uniform blocks (Fig. 3B. *p* = 0.014, Wilcoxon rank-sum test). There were no significant differences in the correlation during the stable phases of normal and uniform blocks (Fig. 3B, *p* = 0.78, Wilcoxon rank-sum test). Thus, pupil responses indicated greater arousal to rare prediction errors during learning.

**Figure 3.**
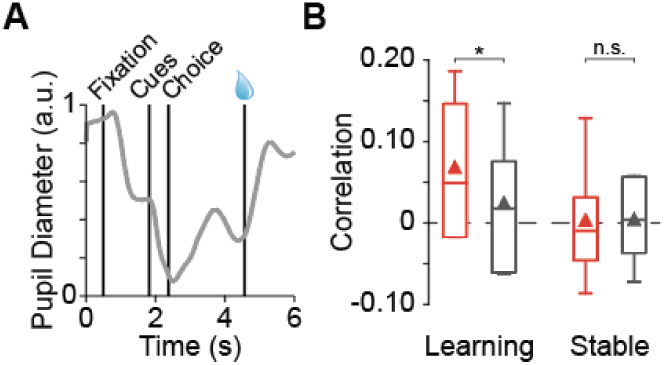
Pupil diameter was correlated with reward prediction error during learning normal distributions. **A.** Averaged normalized pupil diameter response (light grey) to different epochs during behavior. **B.** Box and whisker plots showing the correlation between pupil diameter and RPE, in the learning and stable phases of normal (red) and uniform (grey) blocks. The plot follows the same format as Figure 2B.

### Dopamine responses to rare rewards

After showing that probability distributions were reflected in learning behavior and autonomic responses, we investigated the neural correlates of probability distributions. To do so, we recorded extracellular dopamine neuron action potentials in a passive viewing task wherein monkeys viewed distribution-predicting cues and received rewards (Supplementary Methods). Dopamine neurons were similarly activated by both distribution predicting cues, as would be expected from responses to cues that had similar values (Fig. 4A). At the time of reward delivery, dopamine neurons are activated or suppressed by rewards that are better or worse than predicted, respectively. Therefore, we expected dopamine neurons to be activated by delivery of 0.6 ml and suppressed by delivery of 0.2 ml. Surprisingly, dopamine activations to delivery of 0.6 ml were larger in normal distribution trials, compared to dopamine activations following delivery of the same volume reward in uniform distribution trials (Fig 4B, solid lines). Similarly, dopamine responses were more strongly suppressed by delivery of 0.2 ml reward during normal distribution trials, compared to response suppressions to the same reward during uniform distribution trials (Fig. 4B, dashed lines). As the reward predicting cues had the same subjective values, this response amplification occurred despite the fact that the conventional reward prediction errors were identical (Eq. 1). Moreover, because the amplification was bi-directional – both activations and suppressions were amplified in single neurons – this effect could not be attributed to differences in predicted subjective values. Thus, the effects we observed were robust to errors in the measurement of subjective value.

**Figure 4.**
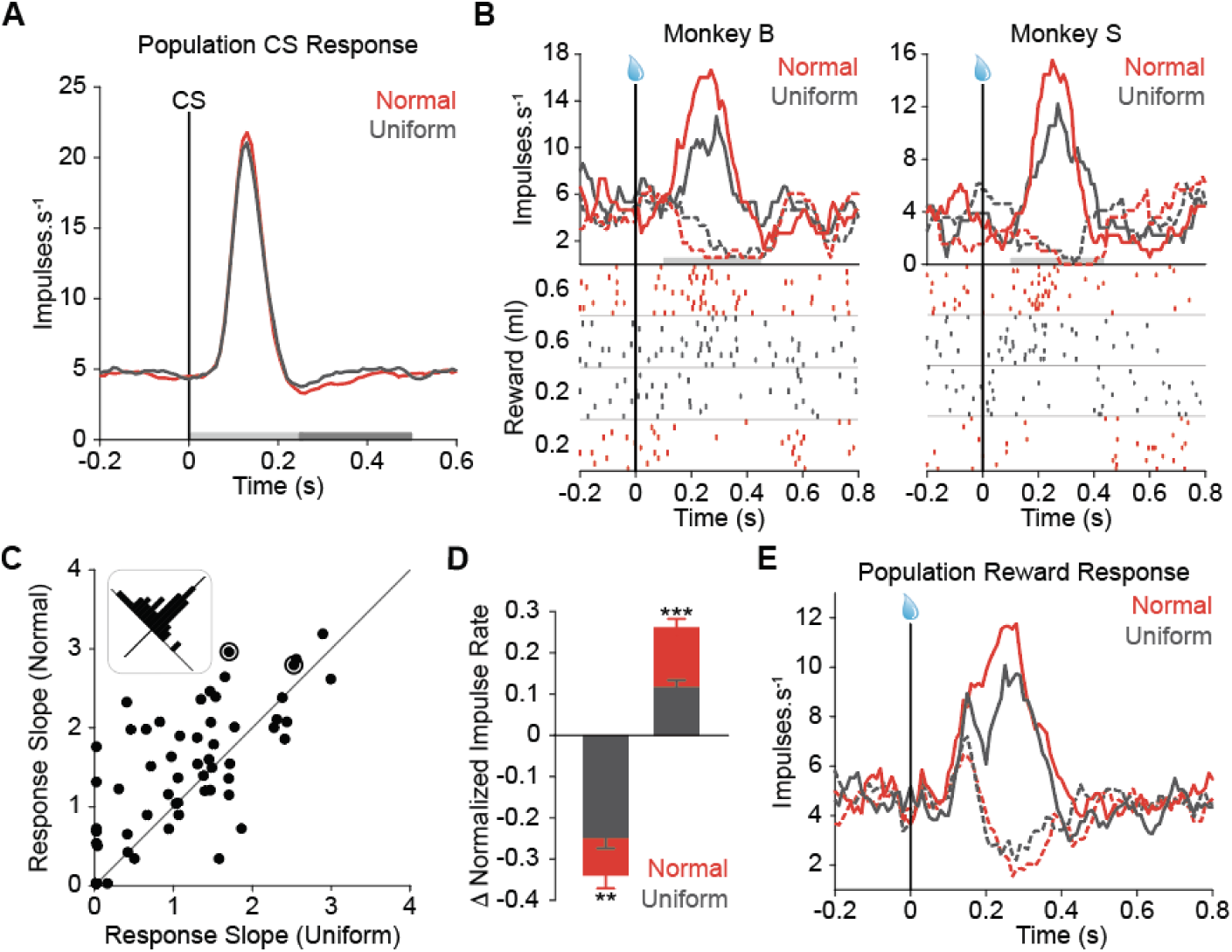
Amplified dopamine reward prediction error responses to rare rewards. **A.** Population PSTHs of conditioned stimuli (CS) responses to the normal and uniform distribution predicting cues. There was no significant difference between the population responses (*p* > 0.6 for both the early and late response components, indicated on the x-axis by the light and dark grey bars, respectively). **B.** Single neuron reward responses from both monkeys. Top: PSTHs show impulse rate as a function of time, aligned to the time of reward (black vertical bar). Light grey bars along with x-axis indicates the response window used for analysis. Bottom: Raster plots, separated by normal and uniform distributions and by reward sizes 0.2 and 0.6 ml. Every line represents the time of an action potential, and every row represents a trial. **C.** Reward response slopes for every neuron in normal and uniform distribution trials. Each dot represents one neuron (*n* = 60 neurons), and the two circled neurons are the example neurons in B. Inset shows the data as a function of the difference from the unity line (diagonal). **D.** Change in normalized impulse rate from baseline in normal and uniform distribution trials, for negative RPE responses (left), and positive RPE responses (right). Error bars are ± standard error of the mean (SEM) across 39 neurons above the unity line. **E.** Population PSTH for neurons that showed amplified responses to the same rewards in the normal and uniform distribution trials. In panels A, B, D, and E, red and grey colors indicate normal and uniform distributions, respectively. In A, B, and E, solid PSTHs represent positive prediction error responses, and dashed PSTHs represent negative prediction error responses.

We sought to quantify the response amplification (Fig. 4B) across neurons. The dynamic ranges of dopamine responses are larger in the positive domain, compared to the negative domain. Therefore, we transformed the responses across the positive and negative domains onto a linear scale (Supplementary Methods). We then used linear regression to measure the response slopes of each neuron to positive and negative RPEs generated during normal and uniform distribution trials. We plotted the response slopes for each neuron in both conditions (Fig. 4C). A majority of neurons had a steeper slope for rare RPEs generated during normal distribution trials, compared to common RPEs generated during uniform distribution trials (Fig. 4C inset, *p* = 0.001, *n* = 60 neurons, one-sample *t*-test). To ensure that the amplification effect was bi-directional and not driven solely by positive responses, we separately analyzed the positive and negative prediction error responses for the 39 neurons above the unity line in Figure 4C. Across this subpopulation, both activations and suppressions were significantly amplified (Fig. 4D, negative and positive prediction error responses *p* < 0.01 and 0.001, respectively, Wilcoxon rank-sum test). There were no significant differences in the positive or negative responses in the subpopulation below the unity line (*p* > 0.05, n = 21, Wilcoxon rank-sum test). Population peri-stimulus time histograms (PSTHs) for the amplified subpopulation show differences in both the positive and negative response domains (Fig. 5E). Thus, phasic dopamine responses are amplified by rare RPEs. These data demonstrate that RPE responses are sensitive to predicted probability distributions.

**Figure 5.**
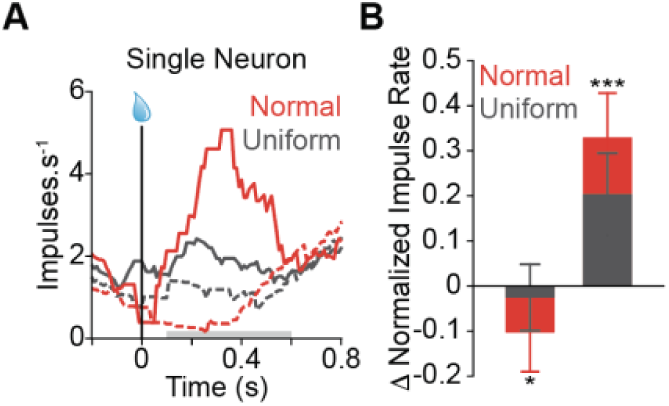
Amplification in dopamine neurons persistent in learning conditions. **A.** Single neuron PSTHs aligned to reward (black vertical line) show amplified responses to positive and negative RPEs in normal distribution trials, compared to uniform distribution trials. Solid PSTHs indicate positive prediction error responses, and dashed PSTHs are negative prediction error responses. The light grey bar along the x-axis indicates the response window used for analysis. **B.** Change in normalized impulse rate from baseline in normal and uniform conditions, for negative RPE responses (left), and positive RPE responses (right). Error bars are ± SEM across neurons. In both panels, red and grey colors indicate normal and uniform distributions, respectively.

The passive viewing task used to collect the neural data (Fig. 4) offers the greatest level of control, because during each trial only one reward-predictive cue is shown to the animals and no choices are required. To validate that the neuronal sensitivity to probability distributions was also present during complex behaviors, we recorded an additional 8 and 12 neurons from monkey B and S, respectively, while the animals learned the predicted values and made choices (Fig. 1C). We used the trial-wise RPE’s derived from the model to categorize each response as a positive or negative prediction error response (Supplementary Methods). We found that positive and negative RPEs elicited amplified neural responses in normal distribution trials, compared to uniform distributions trials (Fig. 5A). The amplification effect was significant in the population (Fig. 5B, negative RPE: *p* = 0.019; positive RPE: *p* = 0.001, *n* = 20 neurons, Wilcoxon rank-sum test). These findings confirm that dopamine responses are sensitive to probability distributions during complex behaviors and demonstrate that amplified dopamine responses can be used to guide active learning and decision making.

## Discussion

Here we show that dopamine-dependent learning behavior and dopamine reward prediction error responses reflect probability distributions. Rare prediction errors – compared to commonly occurring prediction errors of the same magnitude – evoke faster learning, increased autonomic arousal, and amplified neural learning signals. Together, these data reveal a novel computational paradigm for phasic dopamine responses that is distinct from, but complementary to, conventional reward prediction errors. Rare events are often highly significant (*17*). Our data show decision makers exhibit greater vigilance towards rare rewards and learn more from rare reward prediction errors. Amplified dopamine prediction error responses provide a mechanistic account for these behavioral effects.

More than 20 years ago, single unit recordings demonstrated the importance of unpredictability for dopamine neuron responses (*2*). Since then, the successful application of TD learning theory to dopamine signals has largely subsumed the role of unpredictability, and recast it within the framework of value-based prediction errors (*18, 19*). In most experimental paradigms, reward unpredictability is captured by frequentist probability of rewards and, therefore, unpredictability is factored directly into expected value (*5, 7, 20-22*). The resulting Pascalian expected values are learned by TD models (*23*) and reflected in dopamine responses (*5, 20, 22*). Here, we dissociated unpredictability from expected value by pseudorandomly drawing rewards from symmetric probability distributions with equal values. Monkeys learned faster from the more unpredictable rewards drawn from the tails of normal distributions. Likewise, dopamine responses were amplified by greater unpredictability, even when the conventional prediction error described in Eq. 1 was identical. These data reinforce the importance of unpredictability for dopamine responses and learning.

The amplified dopamine responses to rare rewards suggest that reinforcement learning approaches that acquire only the average value of past outcomes are insufficient to describe the phasic activity of dopamine neurons. In fact, dopamine responses distinguish between predicted reward distributions, even when the average expected values and average subjective values of those distributions are identical. Consequently, monkeys and their dopamine neurons appear to learn probability distributions, and this information is used to boost their performance and gain more reward. Therefore, these data suggest that an updated RL algorithm that learns higher moments of probability distributions, such as Kalman TD (*24*), will provide the proper conceptual framework to explain information processing in dopamine responses.

Distribution learning is critical to Bayesian inference and, indeed, the results that we show here are consistent with a signal that could guide Bayesian inference to optimize choices and maximize rewards. However, further experimentation is required to understand whether dopamine signals actually support Bayesian inference. Distribution learning is also emerging as an important approach in machine learning and artificial intelligence (AI) (*25*). Biological learning signals have inspired deep reinforcement learning algorithms with performance that exceeds expert human performance on Atari games, chess, and Go (*26, 27*). The distribution-sensitive neural signals that we observed here could offer further guidance to computational learning; emphasizing the impact of rare events drawn from tails of reward distribution could restrict the necessary search space.

Dopamine reward prediction error signals participate in Hebbian learning (*28, 29*), and these signals are likely responsible for updating action values stored in the striatum (*30*). The amplified dopamine responses coupled with the faster learning dynamics observed here, suggest that the magnitude of dopamine release affected cellular learning mechanisms in the striatum. Moreover, amplified dopamine responses have the ability to modulate dopamine concentrations in the prefrontal cortex (PFC). The level of PFC dopamine is tightly linked to neuronal signaling and working memory performance (*31*). Therefore, amplification of dopamine could explain the exaggerated salience of real – and possibly imagined – rare events, and postulates a neural mechanism to explain aberrant learning observed in mental health disorders, such as schizophrenia.

## Conclusion

Dopamine neurons code reward prediction errors (RPEs), which are classically defined as the differences between received and predicted rewards. However, our data show that predicted probability distributions, rather than just the predicted average values, affects dopamine responses. Specifically, rare rewards drawn from normal distributions amplified dopamine responses, compared to the same rewards drawn more frequently from uniform distributions. Crucially, the classically defined RPEs had identical magnitudes, following rare and common rewards. From a behavioral perspective, rare prediction errors often signal underlying changes in the environment and therefore demand greater vigilance. From a theoretical perspective, this result demonstrates that biologically inspired reinforcement learning algorithms should account for full probability distributions, rather than just point estimates.

## Acknowledgments

The authors would like to thank Andreea Bostan for thoughtful comments and discussion of the manuscript, and Jacquelyn Breter for her work in animal care and enrichment for this project.

## Author contributions

KMR and WRS conceptualized and designed the experiments. KMR, AA, and WRS collected the data. KMR, TH, and WRS analyzed the data. All authors discussed the results of the experiment. KMR, TH, and WRS wrote the manuscript. All authors revised the manuscript.

## Data and materials availability

Available upon request.

## Supplementary Materials

Materials and Methods

Figures S1-S2

References

