## Supplementary Information for "Rare Rewards Amplify Dopamine Learning Responses"

**This PDF file includes:**

Materials and Methods
Figs. S1 to S2
Supplementary References

### Materials and Methods

#### Animals, Surgery and Setup

All animal procedures were approved by Institutional Animal Care and Use Committee of the University of Pittsburgh. We used two male Rhesus macaque monkeys (Macaca mulatta) for these studies (13.9 and 11.2 kg). A titanium head holder (Gray Matter Research) and a recording chamber (Crist Instruments, custom made) were aseptically implanted under general anesthesia before the experiment. The recording chamber for vertical electrode entry was centered 8 mm anterior to the interaural line. During experiments, animals sat in a primate chair (Crist Instruments) positioned 30 cm from a computer monitor. During behavioral training, testing and neuronal recording, eye position was monitored noninvasively using infrared eye tracking (Eyelink Plus 1000). Licking was monitored with an infrared optical sensor positioned in front of the juice spout (Balluff). Eye, lick and digital task event signals were sampled at 2 kHz. Custom-made software (Matlab, Mathworks Inc.) running on a Microsoft Windows 7 computer controlled the behavioral tasks.

#### Behavioral tasks

*Pavlovian task* Three distinct cues (fractal images) were used to predict reward. One predicted a sure reward of 0.4 ml. Another predicted a uniform distribution, where 0.2, 0.4, and 0.6 ml were delivered with equal frequency (1/3 probability for each reward). A final cue predicted a normal reward distribution, where 0.2 and 0.6 ml were delivered with low frequency (2/15 probability for each of the two rewards), and the middle reward (0.4 ml), was delivered with a much higher frequency (11/15 probability). Finally, there was an unpredicted reward condition, where 0.4 ml of juice would be given after no cue was presented. In each trial, one of the three cues, or no cue, was pseudorandomly chosen and was presented to the animal. The reward was delivered 2 s after the cue onset. At the same time, and to assist in reward size identification, a 'value bar cue' (See choice task 1, below) was displayed on the screen that indicated the reward volume. Trials were separated with inter-trial intervals of 2-5 s, chosen from a truncated exponential distribution. Before recording, all cues were well learned after experiencing them repeatedly over multiple sessions (Monkey B: 10 sessions, ~2800 trials; Monkey S: 6 sessions. ~2600 trials).

*Choice task for measuring distribution value.* The monkeys were offered a choice between the well-learned fractal images that predicted a distribution of rewards (Fig. 1A, normal or uniform), and a randomly selected 'safe' alternative value, represented by a value bar cue. The value bar cue had a value range of 0 ml to 0.8 ml, in 0.1 ml increments. Wherever the horizontal bar intersected the vertical scale indicated with 100% certainty the size of juice the monkeys would receive if they chose it. The task was organized so that every session used either the normal distribution cue versus safe values, or the uniform distribution cue versus safe values. The mean of the distribution predicting cue was either 0.3, 0.4, or 0.5 ml. In each choice trial, after successful central fixation for 0.5 s, the two choice options appeared on the monitor and the animal indicated its choice by a saccade towards one of the cues. The animal was allowed to saccade as soon as it wanted. The animal had to keep its gaze on the chosen cue for 0.5 s to confirm its choice. Reward was delivered 1.5 s later. Trials were separated with inter-trial interval of 1.5-6.5 s, drawn from a truncated exponential distribution. Failure to maintain the central fixation or early break of the fixation on the chosen option resulted in a 4 s time-out, and a repeat of the failed trial.

*Choice task to measure learning:* Monkeys were offered two never-before-seen cues in every block of trials. The block length was selected from a truncated exponential distribution between 15 to 25. Each block had both the novel cues drawing rewards from a normal or uniform distribution. Further, each novel cue had a different randomly selected mean that was either 0.2, 0.3, 0.4, 0.5, or 0.6. For example, if it were a uniform block, and the means selected for the two cues were 0.3 and 0.6 ml, the rewards for one cue would be 0.2, 0.3, and 0.4 ml (drawn with equal frequency), and 0.5, 0.6, and 0.7 ml (also drawn with equal frequency). At the start of each new block, the monkeys had to learn which cue was better by sampling the new cues, and estimating which had the higher average reward. In each choice trial, after successful central fixation for 0.5 s, the two choice options appeared on the monitor and the animal indicated its choice by a saccade towards one of the cues. The animal was allowed to saccade as soon as it wanted. The animal had to keep its gaze on the chosen cue for 0.5 s to confirm its choice. Reward was delivered 1.5 s later. Trials were separated with inter-trial interval of 1.5-6.5 s, drawn from a truncated exponential distribution. Failure to maintain the central fixation or early break of the fixation on the chosen option resulted in a 4 s time-out, and a repeat of the failed trial.

**Reinforcement learning models**

_Reinforcement learning models for choice task 2_ We constructed two reinforcement learning (RL) models to examine animals' choices during learning and to acquire trial-by-trial estimate of chosen and unchosen values.

The models had two value functions ($V_{trial}(pd1)$ and $V_{trial}(pd2)$) representing the learned values of probability distribution 1 and probability distribution 2, respectively. In each trial, the probability that the model chooses $pd1$ over $pd2$ was estimated by the softmax rule (*1*) as follows:

$$p\ choose(pd1)_{trial} = \frac{e^{V_{trial}(pd1)/\beta}}{e^{V_{trial}(pd1)/\beta} + e^{V_{trial}(pd2)/\beta}}$$

(Eq. 1)

where $\beta$, the temperature parameter of the softmax rule, determines the level of choice randomness.

In each trial, upon making a choice and receiving an outcome, the value of the chosen option on that trial, $V_{trial}$, was updated according the reward prediction error, as follows:

$$V_{trial+1} = V_{trial} + \alpha_{trial} * \delta_{trial}$$

(Eq. 2)

where $\alpha$ denotes the learning rate, and the prediction error, $\delta_{trial} = r_{trial} - V_{trial}$, indicates the difference between the predicted and realized reward sizes, $V_{trial}$ and $r_{trial}$, respectively. We considered two models to perform the learning rate update. In Model 1, we adopted the dynamic learning rate equation in the Pearce-Hall model as used in (*2*):

$$\alpha_{trial+1} = \eta * |\mathrm{PE}| + (1 - \eta) * \alpha_t$$

(Eq. 3)

In Model 2, we simply asserted the learning rate to be a random walk process, and we reset the learning rate at the start of each block to accommodate our task structure:

$$\begin{cases} \alpha_{trial+1} = \alpha_t + \varepsilon & \text{if t} \neq \text{the end trial of the block} \\ \alpha_{trial+1} = \alpha_m & \text{if t = the end trial of the block} \end{cases}$$

(Eq. 4)

where $\varepsilon$ is Gaussian noise, and $\alpha_m$ is the maximum likelihood estimate, or the initial value, from Model 1.

After fitting the RL models with a forward-filter-backward-smooth algorithm, we used Bayes factor to compare the fitting of two different models on behavioral choices. The less parametrized model, Model 2, was the winning model, with a log likelihood of -1193, where Model 1 had a log likelihood of -1213. Prediction errors (PEs) are then calculated by subtracting the derived value estimates from the rewards monkeys receive. Afterwards, PE is treated as a function of the number of trials since the block shifts.

To characterize learning and stable phases, we used a logarithmic fit of the estimated PEs from the RL model, as the fitted PEs approached 0, the better the animal had learned to estimate the value of the distribution. We also used the absolute size of the PE. If the PE was 0.1, we could assume they had learned the value of the cue – since the true value difference between the lowest and highest values from the mean was 0.1 ml. A change in PE of less than 0.08 from one trial to the next indicated that PE was stable. When both thresholds were met, we consider that the monkeys had entered the stable, asymptotic phase. The transition trials that marked the change from learning to stable were collected block by block, and a Mann-Whitney U test was run to compare the ending positions for normal distribution block and uniform distribution block. We combined data from both animals to have 261 blocks for analysis (Monkey B $n = 178$ blocks, Monkey S $n = 83$ blocks). Separately, both animals took significantly fewer trials in the learning phase for normal versus uniform blocks (Monkey B, $p < 0.001$, Monkey S, $p < 0.001$; Mann-Whitney U test). The faster learning exhibited in the normal distribution block was robust under a wide range of prescribed thresholds.

**Deconvolution**

Event-related pupil responses were analyzed using nideconv, a Python package that specializes in fMRI and pupil signal deconvolution. The design matrix consists of in total twelve event types. There are four pre-reward events for each reward distribution: the onset of central dot for fixation, the onset of cue presentation, the monkeys' saccades to indicate choice and the offset of cue presentation (in temporal order). Based on the Learning/Stable learning period distinction, there are two events put in place to understand the effect of the presence of juice as reward for each reward distribution. The pupil diameter changes related to fixation, the offset of cue presentation, and the presence of rewards are all analyzed 0.5 s pre-event until 2 s post-event. The time window for the onset of cue presentation and monkeys' saccades lasts from 0.5 s pre-event to 3 s post-event. To understand the relationship between pupil diameter and prediction error post-reward, standardized prediction errors and value estimates derived from the Model 2 are first grouped based on reward distribution and learning periods combination, and then are used as covariates in the deconvolution algorithm. Data were analyzed session-by-session. Consequently, we obtain a measure of how sensitive the post-reward pupil diameter changes are to the prediction errors in each reward distribution, by looking at the regression coefficients in the prescribed time window. Given the relatively small variance (mean = 537 ms, standard deviation = 86 ms) in decision time, it did not affect the deconvolved signal.

**Neuronal data acquisition and analysis of neuronal data**

Custom-made, movable, glass-insulated, platinum-plated tungsten microelectrodes were positioned inside a stainless-steel guide cannula and advanced by an oil-driven micromanipulator (Narishige). Action potentials from single neurons were amplified, filtered (band-pass 100 Hz to 3 kHz), and converted into digital pulses when passing an adjustable time–amplitude threshold (Bak Electronics). We stored both analog and digitized data on a computer using custom-made data collection software (Matlab).

Dopamine neurons were functionally localized with respect to (a) the trigeminal somatosensory thalamus explored in awake animals (very small perioral and intraoral receptive fields, high proportion of tonic responses, 2-3 mm dorsoventral extent) (*3*), (b) tonically active position coding ocular motor neurons and (c) phasically direction coding ocular premotor neurons in awake animals. Individual dopamine neurons were identified using established criteria of long waveform (> 2.5 ms, Fig. S1A) and low baseline firing (< 8 impulses/s) (*4*). Following standard sample sizes used in studies investigating neuronal responses in non-human primates, we recorded extracellular activity from 123 dopamine neurons in two monkeys (62 and 61 neurons in Monkeys B and S, respectively). Fifty-four neurons and forty-nine neurons were recorded in a passive viewing task in Monkeys B and S, respectively. Thirty neurons from each animal had a sufficient number of trials (60 total). We used these for further analysis of the Pavlovian data. In the two-alternative forced-choice task we recorded 8 neurons in Monkey B and 12 neurons in Monkey S. We used these for further analysis of the two-alternative forced-choice task data.

The neurons that met these criteria showed the typical phasic activation after unexpected reward, which we used as a fourth criterion for inclusion in data analysis (Fig. S2A, $p < 0.0001$, $n = 80$ neurons; Wilcoxon rank-sum test). Figure S1B and S1C show maps of our recording locations relative to both monkeys' grids, and the number of cells recorded at each location. Figure S1D and S1E show MRI images of Monkey S and the location of the recordings.

We constructed peri-stimulus time histograms (PSTHs) by aligning the neuronal impulses to task events and then averaging across multiple trials. The impulse rates were calculated in non-overlapping time bins of 10 ms. PSTHs were smoothed using a moving average of 70 ms for display purposes. The analysis of neuronal data used defined time windows, individual to each neuron, that included the major positive and negative response components following cue onset and juice delivery, as detailed for each analysis and each figure caption. To compare across a population of neurons, analysis windows were normalized within neurons. Neurons from both monkeys were analyzed together as one population. Response slopes for normal versus uniform were created by individually normalizing negative and positive reward prediction error (RPE) responses from baseline within each neuron to create a linear relationship between negative and positive RPE responses. This was necessary in order to avoid overweighting the positive prediction error in response slope due to the floor effect of negative RPE responses.

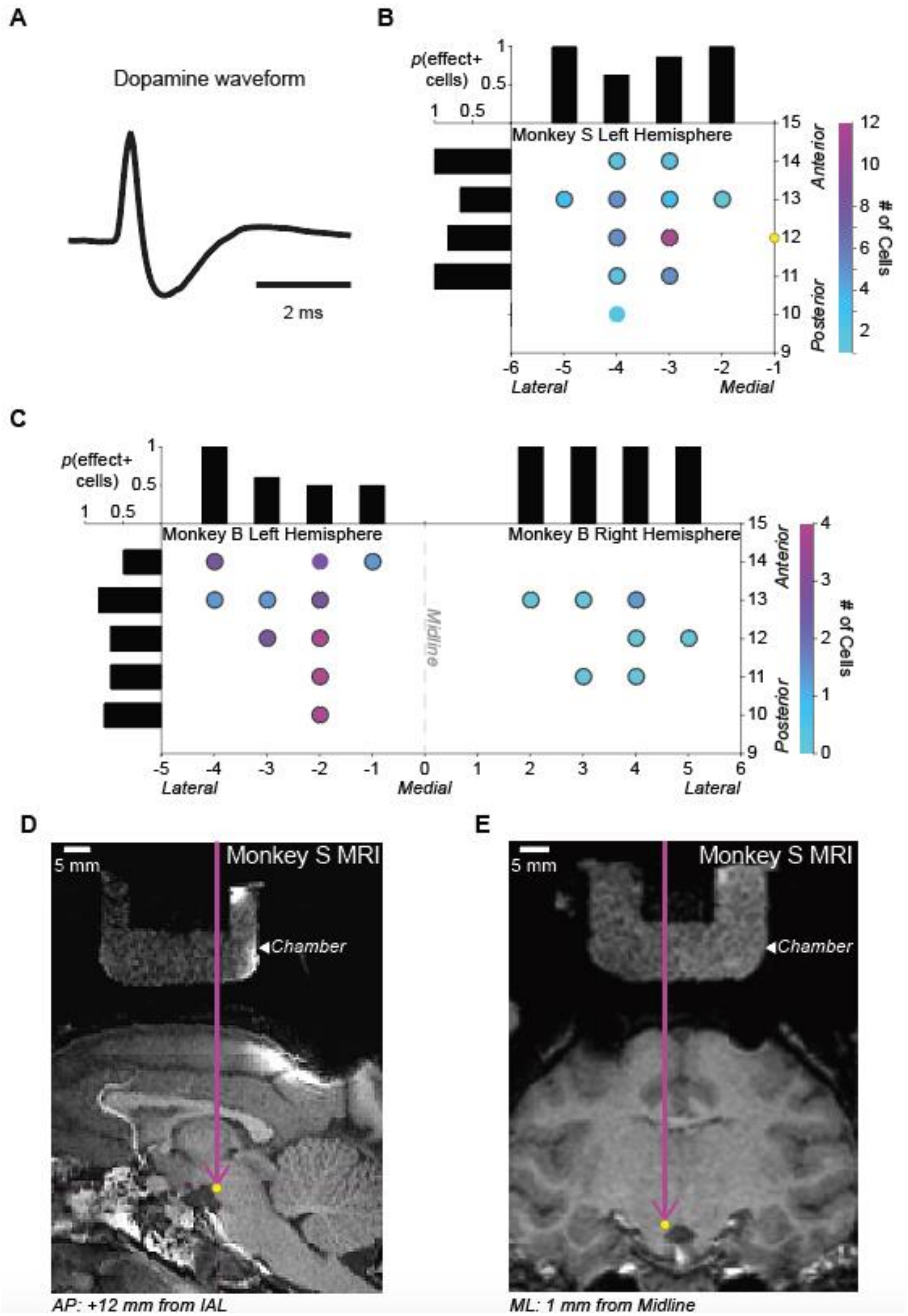


**Figure S1. Dopamine waveform and recording sites. A.** Example dopamine waveform. **B.** Recording locations for the left hemisphere of Monkey S. X-axis indicates lateral to medial location in the grid in millimeters, relative to midline (0). Right y-axis indicates posterior to anterior location in the grid in millimeters, relative to interaural line (IAL). Each locations' color indicates the number of neurons recorded for that location. Black circles surrounding the individual locations indicated that neurons recorded there were part of the population of 39 neurons that had a steeper response slopes in normal compared to uniform condition. Bar graphs on the left and top axes indicate the proportion of cells in that AP (left) or ML (top) location that were effect positive. Yellow dot corresponds to location indicated in MRI scan shown in D and E. **C.** Recording locations for both hemispheres of Monkey B. Same as B. **D.** Sagittal view MRI of the recording chamber of Monkey S. Purple arrow indicates the AP location in the grid (+12 mm from IAL). **E.** Coronal view MRI of the recording chamber of Monkey S. Purple arrow indicates the ML location in the grid (1 mm from Midline). Yellow dot in D and E correspond to approximate recording grid location in B.

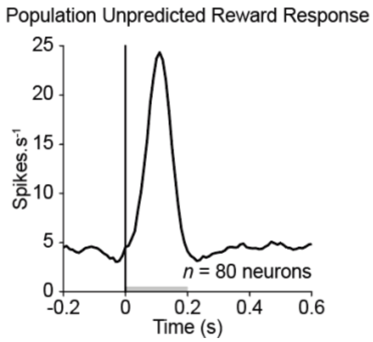


**Figure S2. Unpredicted reward responses for population of dopamine neurons.** The population of 80 neurons used for our analyses in the Pavlovian and choice task had a significant activation following unpredicted reward – a characteristic feature of dopamine neurons. Grey bar along the x-axis indicate the response window used for analysis.
